# Single-trial dynamics of competing reach plans in the human motor periphery

**DOI:** 10.1101/765180

**Authors:** Luc P. J. Selen, Brian D. Corneil, W. Pieter Medendorp

**Author notes:** Corresponding Author: Dr. Luc Selen.

## Abstract

Contemporary motor control theories propose competition between multiple motor plans before the winning command is executed. While most competitions are completed prior to movement onset, movements are often initiated before the competition has been resolved. An example of this is saccadic averaging, wherein the eyes land at an intermediate location between two visual targets. Behavioral and neurophysiological signatures of competing motor commands have also been reported for reaching movements, but debate remains about whether such signatures attest to an unresolved competition, arise from averaging across many trials, or reflect a strategy to optimize behavior given task constraints. Here, we recorded electromyographic activity from an upper limb muscle (m. pectoralis) while twelve (8 female) participants performed an immediate response reach task, freely choosing between one of two identical and suddenly presented visual targets. On each trial, muscle recruitment showed two distinct phases of directionally-tuned activity. In the first wave, time-locked ~100 ms of target presentation, muscle activity was clearly influenced by the non-chosen target, reflecting a competition between reach commands that was biased in favor of the ultimately chosen target. This resulted in an initial movement intermediate between the two targets. In contrast, the second wave, time-locked to voluntary reach onset, was not biased toward the non-chosen target, showing that the competition between targets was resolved. Instead, this wave of activity compensated for the averaging induced by the first wave. Thus, single-trial analysis reveals an evolution in how the non-chosen target differentially influences the first and second wave of muscle activity.

**SIGNIFICANCE STATEMENT:** Contemporary theories of motor control suggest that multiple motor plans compete for selection before the winning command is executed. Evidence for this is found in intermediate reach movements towards two potential target locations, but recent findings have challenged this notion by arguing that intermediate reaching movements reflect an optimal response strategy. By examining upper limb muscle recruitment during a free-choice reach task, we show early recruitment of a sub-optimal averaged motor command to the two targets that subsequently transitions to a single motor command that compensates for the initially averaged motor command. Recording limb muscle activity permits single-trial resolution of the dynamic influence of the non-chosen target through time.

## INTRODUCTION

The environment offers multiple action opportunities, but ultimately only one action can be selected. Classic decision-making theories assume a two-stage process, where the brain selects an appropriate action, and then plans and executes the desired motor commands (Donders, 1969; McClelland, 1979). However, neurophysiological studies have suggested that multiple potential motor plans can be concurrently encoded and compete for selection within brain regions involved in eye (Christopoulos et al., 2018; McPeek and Keller, 2004; Port and Wurtz, 2003) or reach movements (Cisek and Kalaska, 2005; Klaes et al., 2011). Competition may also influence behavioral output. For example, when free to look to either one of two suddenly appearing visual targets, participants sometimes look to an in-between position (Chou et al., 1999; Findlay, 1982). Because such saccadic averaging is most prominent for short-latency saccades (Ottes et al., 1984; Walker et al., 1997), it is thought that target representations initially compete for selection, before resolving into a final decision (Kim and Basso, 2008; McPeek and Keller, 2002).

There is currently debate on whether reach motor plans can also be represented concurrently. Recent neurophysiological results (Dekleva et al., 2018) suggest that apparent concurrent encoding of multiple reach plans (Cisek and Kalaska,2005) may arise from averaging neural activity across many trials; while the represented alternative can vary trial-to-trial, only one alternative is represented on any given trial. Reach trajectories intermediate between two alternatives have been observed in ‘go-before-you-know’ paradigms (Chapman et al., 2010), in which reach movements start before the ‘correct’ target is unveiled. Such intermediate reaches have been ascribed to averaging of competing reach plans (Gallivan et al., 2017; Stewart et al., 2014; Enachescu et al., 2021), or to strategic optimization of success given task constraints (Haith et al., 2015; Hudson et al., 2007; Wong and Haith, 2017; Alhussein and Smith, 2021).

For reaching, target competition studies often impose a delay between target presentation and movement initiation or target identification (as in the ‘go-before-you-know’ paradigm). This approach differs from the immediate and free response paradigms that elicit saccadic averaging. Here, we employ an immediate response paradigm to show competition between potential reach targets at the individual trial level, studying humans reaching in a *free-choice, double-target task* (**Fig. 1b,c**). Unlike the ‘go-before-you-know’ paradigm, there is no correct target, and nothing is gained by strategically aiming between the two targets. We recorded electromyographic (EMG) activity (m. pectoralis) and analyzed timing and magnitude of recruitment in response to target presentation.

**Figure 1:**
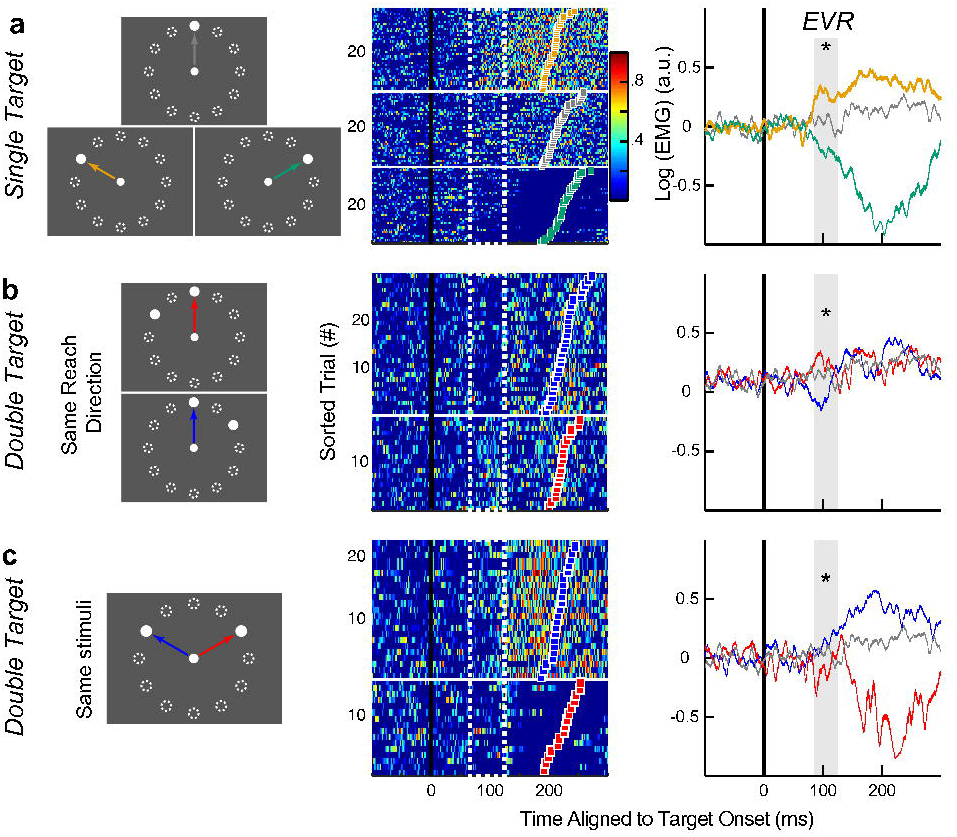
EVR from a representative participant. **A.** Individual (middle panel) and mean ± SEM (right) log-normalized EMG-activity from the right PEC muscle during left-outward (yellow), straight outward (gray), and right-outward (green) *Single Target* reach trials (left panel). All EMG-activity is aligned to the onset of the peripheral visual target (thick black vertical line). For the middle panel, each row represents EMG-activity within a single trial and trials were sorted based on reach RT (colored squares). Dashed white box and shaded area in the individual and mean EMG plots represent the EVR-epoch (85-125 ms after stimulus onset). **B.** EMG-activity for *Double Target* trials when matched for the same outward reach movement. The non-chosen target was either 60° CW (blue) or CCW (red) of the reach target. Same layout as **A**. **C.** EMG-activity for 120°*Double Target* trials for the same visual target layout, but different chosen target directions. Same layout as **A**. * *P* < 0.05.

For reaches to a single visual target, two waves of directionally-tuned EMG-activity have been observed (Glover and Baker, 2019; Pruszynski et al., 2010; Wood et al., 2015): an initial stimulus-locked response ~100 ms after visual target onset, which we refer to as an express visuomotor response (EVR, Contemori et al., 2021a), followed by a larger wave of EMG-activity predictive of the onset of the reach movement (MOV, ~200-300 ms after the EVR). Muscle recruitment during the EVR and MOV-interval are governed by different processes. For example, the EVR is directed towards the stimulus location even during anti-reaches (Gu et al., 2016), is only influenced by the implicit but not explicit component of motor learning (Gu et al., 2019), and depends on stimulus properties (Glover and Baker, 2019; Kozak et al., 2019; Wood et al., 2015; Kozak and Corneil, 2021) and cueing (Contemori et al., 2021b). In contrast, MOV-epoch activity reflects the actual reach kinematics, but is not influenced by stimulus properties.

Examining EMG-activity in these intervals during free choice, double-target reaching, suggests a dynamically evolving interaction between the chosen and non-chosen target. During the EVR-interval, the non-chosen target influences muscle recruitment, revealing averaging that is biased in favor of the ultimately selected target. This averaging produces a subtle attraction of early reach kinematics to the non-chosen target. Subsequently, this non-chosen target influence yields into a goal-directed motor command during the MOV-interval, compensating for the earlier subtle attraction by bowing the reach trajectory away from the non-chosen target.

## MATERIALS AND METHODS

### Participants and Procedures

The experiment was conducted with approval from the institutional ethics committee from the Faculty of Social Sciences at Radboud University Nijmegen, The Netherlands. Twelve participants (eight females and four males), between 18 and 33 years of age (mean ±SD = 24 ± 5), gave their written consent prior to participating in the experiment. Three participants (one female and two males) were self-declared left-handed, while the remaining participants were self-declared right-handed. All participants were compensated for their time with either course credits or a monetary payment and they were free to withdraw from the experiment at any time. All participants had normal or corrected-to-normal vision and had no known motor impairments.

### Reach Apparatus and Kinematic Acquisition

Participants were seated in a chair in front of a robotic rig. The participant’s right arm was supported by an air-sled floating on top of a glass table. All participants performed right-handed horizontal planar reaching movements while holding the handle of the planar robotic manipulandum (vBOT, Howard et al., 2009). The vBOT measured both the *x-* and *y*-positions of the handle at a 1 kHz sampling rate. Throughout the whole experiment a constant load of 5 N in the rightward direction, relative to the participant, was applied to increase the baseline activity for the right pectoralis muscle (see below). All visual stimuli were presented within the plane of the horizontal reach movements via a mirror, which reflected the display of a downward facing LCD monitor (Asus – model VG278H, Taipei, Taiwan). The start position and the peripheral visual targets were presented as white circles (0.5 and 1.0 cm radii, respectively) onto a black background. Real-time visual feedback of the participant’s hand position was given throughout the experiment and was represented by a yellow cursor (0.25 cm in radius). Vision of the physical arm, hand and manipulandum was occluded by the mirror.

### EMG Acquisition

EMG-activity was recorded from the clavicular head of the right pectoralis major (PEC) muscle using wireless surface EMG electrodes (Trigno sensors, Delsys Inc., Natick, MA, USA). The electrodes were placed ~1 cm inferior to the inflection point of the participant’s right clavicle. Concurrent with the EMG recordings, we also recorded a photodiode signal that indicated the precise onset of the peripheral visual targets on the LCD screen. Both the EMG and photodiode signals were digitized and sampled at 1.11 kHz.

### Experimental Paradigm

Each trial began with the onset of the start position located at the center of the screen, which was also aligned with the participant’s midline. Participants had to move their cursor into the start position and after a randomized delay period (1-1.5 s) either one (*Single Target*, 25% of all trials, **Fig. 1a**) or two peripheral targets appeared (*Double Targets*, 75%, **Fig. 1b,c**). All peripheral targets were presented 10 cm away from the start position and at one of 12 equally spaced locations around the start position (dotted circles in **Fig. 1a**). The onset of the peripheral targets occurred concurrently with the offset of the start position. Participants were explicitly instructed to reach as fast as possible towards one of the peripheral target locations during *Double Target* trials. To ensure that the participants reached as fast as possible, the peripheral targets turned red if the cursor had not moved out of the start position within 500 ms after the onset of the peripheral targets. The trial ended as soon as the cursor entered one of the peripheral targets. Note that it is highly unlikely and suboptimal for participants to make anticipatory movements, given the high degree of spatial uncertainty of where targets would appear across trials.

For every participant, the experiment consisted of eight blocks, each block contained 240 trials, with 60 *Single Target* and 180 *Double Target* trials, pseudo-randomly interleaved. For the *Double Target* trials, the two targets appeared either 60°, 120°, or 180° apart in equal likelihood. Each possible single and double target configuration was presented five times in every block, resulting in 40 repeats over the whole experiment. This design is expected to average out any trial history effects.

### Data Analyses

All data were analyzed using custom-written scripts in Matlab (version R2014b, Mathworks Inc., Natick, MA, USA). For both the 60° and 120°*Double Target* trials, we sorted trials based on whether the final reach was directed to either the CW (**Fig. 1b**, red arrow) or CCW target location (blue arrow). Thus, for all CW and CCW reach trials the non-chosen target location was in the CCW and CW direction, respectively. Trials from the 180°*Double Target* condition cannot be sorted in this way, since the non-chosen target location was always 180° away.

#### Reach onset detection and initial reach error

Reach onset was identified as the first time-point after the onset of the peripheral targets at which the hand speed exceeded 2 cm/s. Reach reaction time (RT) was calculated as the time between the onset of the peripheral targets and the initiation of the reach movement. Initial reach direction was quantified as the angular direction of the vector between the start position and the location of the hand at the time of reach onset. From this, initial reach error was defined as the angular difference between the initial reach direction and the direction of the final chosen target.

#### EMG processing and trial inclusion criteria

All EMG data were first rectified and aligned to both the onset of the peripheral targets (as measured by the onset of the photodiode) and reach initiation. To account for the difference in EMG recordings across the participants, we first normalized EMG-activity for each participant by dividing against their own mean baseline activity (i.e. mean EMG-activity over the 40 ms window prior to the stimuli onset). We then log-normalized each participants EMG-activity to account for the non-linearity of EMG-activity. This normalization transforms the distribution of EMG values from an exponential distribution, with many values close to zero and few large values, into a normal distribution. We specifically examined two distinct epochs of EMG-activity: (1) The initial EVR, that is evoked 85-125 ms after the onset of the visual stimuli (Gu et al., 2016, 2018, 2019), and (2) the later movement-related response (MOV, −20 to 20 ms around reach initiation) associated with reach onset. To prevent any overlap between these two different epochs (Gu et al., 2016, 2019; Kozak et al., 2019), we excluded all trials with RTs less than 185 ms (~7% all trials). We also excluded with RTs greater than 500 ms (<0.1% of all trials).

#### Receiver-Operating Characteristic Analysis

As done previously (Corneil et al., 2004; Pruszynski et al., 2010), we used a time-series receiver-operating characteristic (ROC) analysis to quantitatively detect the presence of an EVR. To do this, we first separated leftward (target locations between 120° and 240° from straight right) and rightward (−60° to 60°) *Single Target* trials. For each timepoint from 100 ms before to 300 ms after target onset, we calculated the area under the ROC curve between the EMG-activity for leftward compared to rightward trials. This metric indicates the probability that an ideal observer could discriminate the target location based solely on the distribution of EMG-activity at that given timepoint. A value of 0.5 indicates chance discrimination, whereas a value of 1 or 0 indicates perfect correct or incorrect discrimination, respectively. We set the threshold for discrimination at 0.6, as this criterion exceeds the 95% confidence intervals for EMG data that has been randomly shuffled through a bootstrapping procedure (Chapman and Corneil, 2011). The discrimination time was defined as the first timepoint after target onset at which the ROC metric was above 0.6 and remained above that threshold for at least five out of the next 10 timepoints. We defined any participant with a discrimination time less than 125 ms as a participant exhibiting a EVR. Based on this criterion, 11 of the 12 participants had a detectable EVR. All subsequent analyses were done on the 11 participants with an EVR.

#### Directional tuning of EMG-activity

We assumed cosine tuning (**Eq. 1**) between the log-normalized EMG-activity and the chosen target location for both the EVR and MOV-epochs:

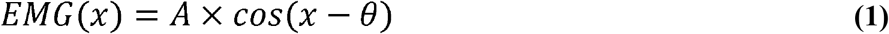

in which *x* is the chosen target location in degrees, starting CCW from straight right; *EMG(x)* is the log-normalized EMG-activity for the given target location; *A* is the amplitude of the cosine tuning; and *θ* is the preferred direction (PD) of the EMG-activity. We used Matlab’s curve fitting toolbox *fit* function to estimate both the *A* and *θ* parameters. We constrained our search parameters such that *A* > 0 and 0° ≤ θ < 360°. The initial search parameters were *A* = 1 and *θ* = 180°. PDs of 0° and 180° would represent straight rightward and leftward, respectively.

#### Model predictions

Previous studies have proposed different models of how the brain converts multiple visual targets into a single motor command. Here we assumed a constant non-linear cosine tuning between target locations and motor commands in *Single Target* trials to generate the predicted responses during *Double Target* trials. Each model used parameters derived from each participant’s own *Single Target* data (**Fig. 2a**) to predict both the PD and amplitude of the cosine tuning curves for *Double Target* trials. Thus, no free parameters were fitted in any of these four models.

**Figure 2:**
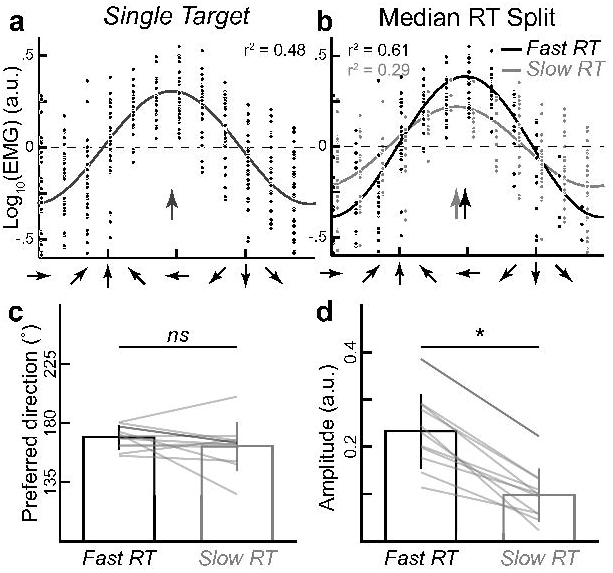
Directional tuning of the EVR during *Single Target* trials. **A.** Cosine tuning of log-normalized EVR magnitude as a function of the target direction for *Single Target* trials from the representative participant in **Figure 1**. Dots indicate each trial, the solid line indicates the fit, and the arrow indicates the PD of the fit. **B.** The cosine tuning is maintained regardless of the ensuing reach RT, same data as **a**. but re-fitted for *Fast* (black) and *Slow RT* (gray) *Single Target* trials separately. For illustration purposes only, we have staggered the individual trial data to illustrate the difference between the two conditions. We did not stagger the cosine tuning curves. **C. D.** Group (*n* = 11) mean ± SEM for the PD (c) and amplitude (**d**) of the fits between the *Fast* and *Slow RT* trials. Each gray line indicates an individual participant, and the darker line indicates the representative participant. * *P* < 0.05.

##### Model 1

The *winner-take-all model* (**Fig. 4a**) assumes that only the target location that the participant reaches towards is converted into a motor command. Therefore, *EMG*(*x*_1_|*x*_1_, *x*_2_) = *EMG*(*x*_1_), where *x*_1_ and *x*_2_ are the chosen and non-chosen target locations, respectively.

**Figure 3:**
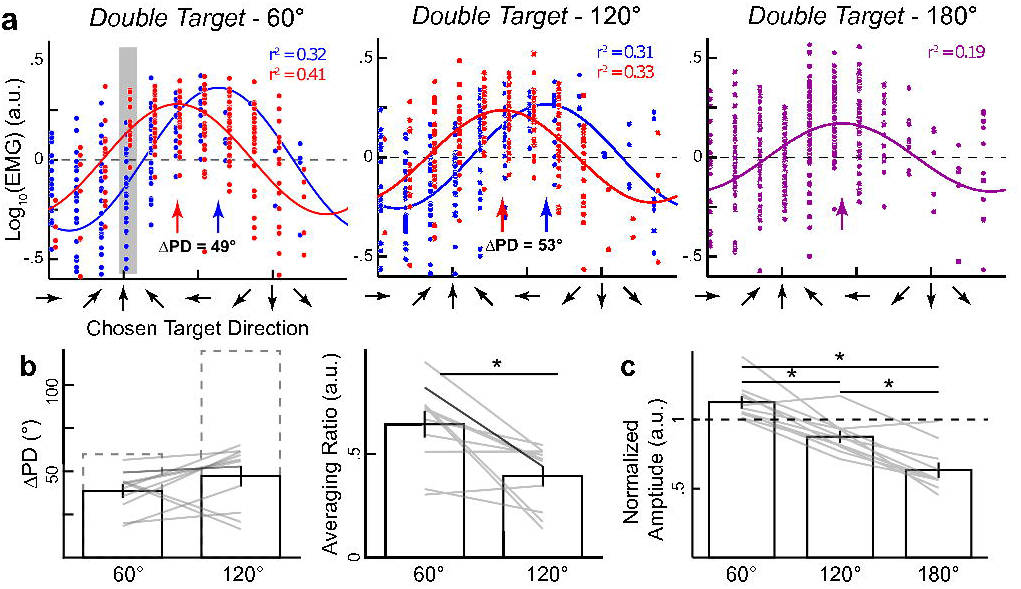
Systematic changes in directional tuning of the EVR during *Double Target* trials. **A.** Fits for 60°, 120° and 180° conditions of the *Double Target* trials with all data aligned to the chosen target direction from the representative participant. For both the 60° and 120° conditions, the trials were sorted based on whether the non-chosen target was either CW (blue) or CCW (red) of the chosen target direction. Data in the shaded panel indicates the trials from **Figure 1b**. **B.** Group mean ± SEM shifts in PD (ΔPD) between the CW and CCW trials (left panel) and the normalized averaging ratio (right) for both 60° and 120° conditions across our participants. Dashed box indicates the predicted ΔPD if the EVR would be a complete average of the two targets (averaging ratio = 1 a.u.). **C.** Mean ± SEM amplitude of the fits for the three different *Double Target* conditions across our participants. The amplitudes were normalized to each participant’s own amplitude fit from the *Single Target* trials. Each grey line indicates a different participant. * *P* < 0.05.

**Figure 4:**
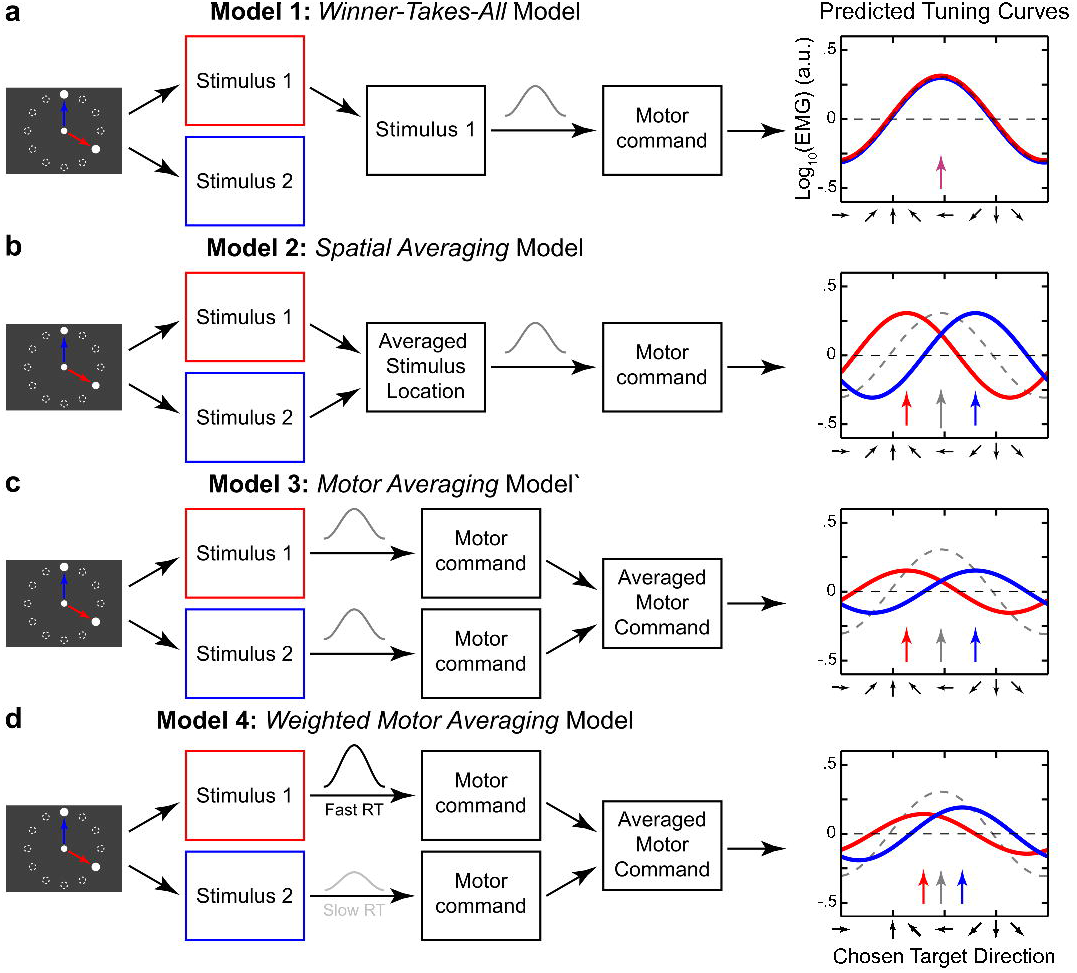
Model predictions of the tuning curves during *Double Target* trials. **A.** The *winner-takes-all* model chooses one visual stimulus as the target and converts it into the final motor command. **B.** The *spatial averaging* model averages the two visual stimulus directions into an intermediate target direction and that target direction is converted into a motor command. **C.** The *motor averaging* model first converts the two visual stimuli into two separated motor commands. Then it averages the two motor commands into a single motor command. **D.** The *weighted motor averaging* model first converts the two visual stimuli into two separate motor commands, but the cosine tuning have different weights. Then it averages the two motor commands into a single motor command. For the right column: red curves for CW chosen target, blue curves for the CCW chosen target and dashed grey curve the single target tuning curve.

##### Model 2

The *spatial averaging model* (**Fig. 4b**) assumes that the two potential target locations are first spatially averaged into an intermediate target location. Then that target location is converted into a motor command. Therefore, 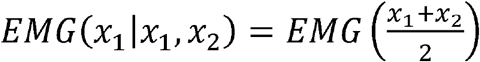.

##### Model 3

The *motor averaging model* (**Fig. 4c**) assumes that the two potential target locations are first converted into their own distinct motor commands and then averaged into a single motor command. Therefore, *EMG*(*x*_1_|*x*_1_, *x*_2_) = 0.5 × *EMG*(*x*_1_) + 0.5 × *EMG*(*x*_2_).

##### Model 4

The *weighted motor averaging model* (**Fig. 4d**) is a variation of the motor averaging model. It assumes that the two target locations are first converted into their associated motor commands, which are then differentially weighted before being averaged into a single motor command. A higher weight is assigned to the chosen target compared to the non-chosen target location. To estimate these weights indirectly, we used each participant’s own *Single Target* data. Previous studies have shown that the EVR magnitude is negatively correlated with the ensuing RT for single target visually-guided reaches (Pruszynski et al., 2010; Gu et al., 2016). We assumed that the trial-by-trial magnitude of the EVR reflected the ‘readiness’ to move towards the target location. Thus, we performed a median RT split of the *Single Target* data to get cosine tuning for both *Fast* and *Slow RT* trials (**Fig. 2a**). This results in *Fast RT* and *Slow RT* amplitude and PD estimates, which were used to compute the tuning curves for the *Double Target* trials. The *Fast RT* and *Slow RT* parameters were used for the chosen and non-chosen target location, respectively. Therefore, *EMG*(*x*_1_|*x*_1_, *x*_2_) = 0.5 × EMG_Fast_ + 0.5 × *EMG_slow_*(*x*_2_).

To quantify the goodness-of-fit for each model, due to the non-linear interaction between PD and normalized amplitude, we evaluated the total fit error between the predicted and observed tuning curves. To do this, we took the sum of mean squared error for each of the 12 different reach directions (i.e. *x*_1_ = 0°, 30°, 60°,… 330°) between the predicted and observed tuning curves.

### Statistical Analyses

Statistical analyses were performed using either one or two-sample *t*-tests or a one-way ANOVA. For all post-hoc comparisons, we used a Tukey’s HSD correction. The statistical significance was set as *P* < 0.05. For the model comparison, significance was set at *P* < 0.0083, Bonferroni corrected for the six possible comparisons between the four different models.

## RESULTS

Under continuous EMG recording of the right PEC muscle, participants performed a free-choice goal-directed center-out right-handed reach movement in response to the onset of either one (**Fig. 1a**, *Single Target*) or two visual targets (**Fig. 1b** and **c**, *Double Target* trials) that appeared concurrently. The visual targets pseudo-randomly appeared at 12 different possible directions equally spaced around the start position. For *Double Target* trials, the two visual stimuli had an angular separation of either 60°, 120°, or 180°. Choice probability for the clockwise or counterclockwise target in *Double Target* did not differ from 0.5 for both the 60° and 120° target separation (*P_CW,60_ = 0.52 ± 0.04; p = 0.06 and P_CW, 120_ = 0.51 ± 0.02; p = 0.13*).

Prior to examining the *Double Target* trials, we will first describe PEC EMG-activity during the *Single Target* trials. **Figure 1a** shows the individual (middle panel) and mean log-normalized EMG-activity (right) during left-outward (orange trace), straight outward (gray), and right-outward *Single Target* trials (green) from a representative participant. All trials are aligned to visual target onset and the individual trials were sorted based on the reach RTs (color squares). Note, the increase and decrease of activity for the right PEC muscle for left-outward and right-outward reach movement, respectively. Consistent with previous studies (Pruszynski et al., 2010; Wood et al., 2015; Glover and Baker, 2019), we observed a reliable difference in EMG-activity for the three different reach directions at two epochs: an initial EVR-epoch that occurs ~100 ms after stimulus onset and a later MOV-epoch associated with reach RT (stochastically occurring ~200 ms after stimulus onset). Across our participants, the mean ±SEM discrimination time (see **Materials and Methods**) for the EVR was 88 ±3 ms and the corresponding reach RT was 232 ± 3 ms. We calculated the EVR magnitude for a given trial as the mean log-normalized EMG-activity during the EVR-epoch, 85-125 ms after stimulus onset (Gu et al., 2018, 2019) indicated by the white dashed boxes and shaded panels in **Figure 1**. For this participant, we found a reliable increase and decrease in EVR magnitude for left-outward and right-outward trials, respectively, when compared to straight outward trials (1-way ANOVA, *F*_(2,105)_ = 37.4, *P* < 10^-12^, post-hoc Tukey’s HSD, both *P* < 0.001).

Having established the profile of EMG-activity during the EVR-epoch on *Single Target* trials, we next examined if the presence of a second non-chosen target during the *Double Target* trials changed the EVR. For a direct comparison with **Figure 1a**, we first examined trials with the same reach direction (i.e. straight outward) but with a different non-chosen target location (60° CW, blue, or 60° CCW, red, from the target, **Figure 1b**). If the non-chosen target location has no influence (i.e. no averaging), we would predict that the EVR magnitude resembles that observed during outward reach movement during *Single Target* trials, which we overlaid in gray in **Figure 1b**. Despite the same reach direction, we observed both an increase and a decrease of EMG-activity during EVR-epoch for *Double Target* trials relative to the *Single Target* trials (1-way ANOVA, *F*_(2,83)_ = 16.2, *P* < 10^-5^, post-hoc Tukey’s HSD, *P* = 0.01 and *P* = 0.004, respectively) when the non-chosen target was in the left-outward and right-outward locations, respectively. This result suggests that EMG-activity during the EVR is systematically altered by the presence of a second non-chosen target.

A second way to examine the EVR during *Double Target* trials is to compare EVR magnitude on trials with the same two visual targets, but different reach directions. **Figure 1c** shows the EMG-activity when both the left-outward (blue) and right-outward (red) targets were presented to the representative participant. If the EVR averaged the locations of the two visual targets completely, then we would predict that the resulting EMG-activity would not differ regardless of the final reach direction. However, we observed a reliable difference in the EVR, with it being slightly larger when the participant chose the left-outward versus right-outward target (1-way ANOVA, *F*_(2,72)_ = 7.06, *P* = 0.002, post-hoc Tukey’s HSD, *P* = 0.01). This result suggests that EMG-activity during the EVR is modulated by the chosen reach direction, even when the same two visual targets are presented.

### Systematic shifts in tuning of the EVR during Double Target trials

The results from **Figure 1b** and **c** demonstrate that the magnitude of the EVR during *Double Target* trials depended on both the target configuration and the eventual reach direction. To quantify the extent of averaging that occurred, we sought to compare how the directional tuning of the EVR changed between *Single* and *Double Target* trials. Previously, it has been shown that the log transformed EVR magnitude can be described by a cosine tuning function (Gu et al., 2019; **Eq. 1**). For each tuning function we can extract both the preferred direction (PD) and the amplitude of the fit. **Figure 2a** shows both individual trial data (dots) and the cosine tuning fit (line) for the *Single Target* trials from the representative participant in **Figure 1a**. The PD of this fit was 173° CCW (arrow) from straight rightward, indicating that the largest EVR magnitude could be evoked by a visual target presented straight leftward of the start position. Importantly, this cosine tuning between EVR magnitude and target location was not simply due to movement-related EMG-activity from trials with the shortest RTs, as this relation was still present when we performed a median RT split and re-fitted the data on either *Fast RT* (**Figure 2b**, dark line) or *Slow RT* trials (light), separately. Across our participants, we found no systematic difference in the PDs between *Fast* and *Slow RT* trials (**Figure 2c**, group mean ±SEM: PD = 169°± 3°and 162° ± 5°, respectively, paired *t*-test, *t*_(10)_ = 1.30, *P* = 0.22). We did find larger amplitudes (i.e. larger EVR magnitudes) for *Fast* compared to *Slow RT* trials (**Figure 2d**, paired *t*-test, *t*_(10)_ = 7.89, *P* < 10^-4^), which is consistent with previous studies demonstrating a negative correlation between EVR magnitudes and RTs on a trial-by-trial basis (Pruszynski et al., 2010; Gu et al., 2016). Note, we will leverage this relationship later in the modeling portion of the **RESULTS**.

We next fitted the EVR cosine tuning for the *Double Target* trials. For this we chose to align the trials based on the participant’s reach direction (**Figure 1b**) rather than controlling for the visual target locations (**Figure 1c**) to accentuate the effect of the non-chosen target location. **Figure 3a** shows the fits for the three different angular separations for the representative participant. For both the 60° and 120° conditions, we generated two separate fits for when the non-chosen target location was either CW (red) or CCW (blue) relative to the reach direction. To give more intuition of how this figure relates to individual trials, the highlighted data (shaded box in the left panel of **Figure 3a**) corresponds to the same trials as **Figure 1b**. The right panel of **Figure 3a** shows the fit of EVR magnitude to the 180°*Double Target* condition. Note, the data cannot be split because the non-chosen target location is always 180° away from the reach direction. Despite the two targets being in diametrically opposite directions, the EVR was still reliably tuned for the 180° condition (r^2^ = 0.19, for this participant). Across participants, the directional tuning of the EVR during the 180° *Double Target* trials was not reliably different from that observed in the *Single Target* trials (paired *t*-test, *t*_(10)_ = 1.92, *P* = 0.08), although we did find a systematic decrease in the amplitude of the fits (see below).

For both the 60° and 120° conditions, since we aligned our data relative to the final reach direction, the only difference between CW and CCW trials was the non-chosen target location. If the EMG-activity was the result of a perfect averaging between the two target locations, then we would predict the difference in PD between CW and CCW trials (ΔPD) to be equal to the angular separation between the two targets (i.e. ΔPD = 60° and 120°, respectively). If the EMG-activity was only influenced by the chosen target direction, then we would predict no difference between CW and CCW conditions (ΔPD = 0°). Consistent with the individual trial data from **Figure 1b**, we observed signs of averaging, albeit incomplete, for the representative participant for both the 60° and 120° conditions, with ΔPDs of 49.3° and 53.0°, respectively (**Figure 3a**).

We found similar results of partial averaging across our participants for both the 60° (**Figure 3b**, left panel, mean ± SEM, ΔPD = 38.6° ± 3.5°, one sample *t*-test against zero, *t*_(10)_ = 10.9, *P* < 10^-6^) and 120° *Double Target* conditions (ΔPD = 47.2° ± 5.4°, one sample *t*-test, *t*_(10)_ = 8.7, *P* < 10^-5^). To fairly compare the extent of averaging between the conditions, we converted the ΔPD into an averaging ratio (**Figure 3b**, right panel): a value of 1 indicates complete averaging (ΔPD = 60° and 120°, dashed lines) and a value of 0 indicates no averaging (ΔPD = 0°). Overall, we found that the extent of averaging decreases as the angular separation increased from 60° to 120° (averaging ratio = 0.6 ± 0.06 and 0.39 ± 0.05 a.u., respectively, paired *t*-test, *t*_(10)_ = 3.81, *P* = 0.003).

In addition to the changes in PD of the EVR tuning, we also quantified the changes in the amplitude during *Double Target* trials. **Figure 3c** shows the mean amplitude for the three conditions, normalized to each participant’s own *Single Target* amplitude as a baseline. We observed a systematic decrease in amplitude as a function of angular separation: 1.13 ± 0.04, 0.88 ± 0.04, and 0.63 ± 0.05 a.u. for the 60°, 120°, and 180° conditions, respectively (repeated measures 1-way ANOVA, *F*_(2,20)_ = 41.1, *P* < 10^-7^, post-hoc paired *t*-test, all *t*_(10)_ > 5.5, *P* < 10^-3^). The systematic changes in PD and amplitude will be interpreted based on different possible averaging models tested below.

### Model predictions of EMG-activity during the EVR-epoch for Double Target trials

Previous studies examining averaging behavior for both eye and reach movements have proposed different models for how the two visual targets may be integrated into a single motor command. These models make distinct predictions for how the PD and amplitude of the tuning curves should change between *Single* and *Double Target* trials (see **Materials and Methods** for exact details). **Figure 4,** right column, shows the predicted tuning curves generated from the four different proposed models for both the 120° CW and CCW conditions, using the *Single Target* data (dashed gray line) from the representative participants. **Model 1** is the *winner-takes-all model* (**Figure 4a**), which proposes that the two visual targets compete for selection in a winner-takes-all process, resulting in a motor command that is generated towards the winning target location (Donders, 1969; McClelland, 1979). Effectively, there is no integration between the two target locations at any stage of the process. Note this model is agnostic about whether the competition for selection occurs at either a spatial or motor representation. **Model 2** is the *spatial averaging model* (**Figure 4b**), which proposes that the two targets are first averaged into a spatial representation, resulting in a motor command towards the intermediate spatial direction (Findlay, 1982; Glimcher and Sparks, 1993; Walker et al., 1997; Chou et al., 1999). **Model 3** is the *motor averaging model* (**Figure 4c**), which proposes that the two targets are first converted into two independent motor commands (Edelman and Keller, 1998; Port and Wurtz, 2003; Cisek and Kalaska, 2005) and then averaged into a single motor command (Katnani and Gandhi, 2011; Stewart et al., 2014; Gallivan et al., 2017). Finally, **Model 4** is the *weighted motor averaging model* (**Figure 4d**), which is a variation of the motor averaging model. Once again, the two targets are first converted into two separate motor commands, but a stronger weighting is given towards the chosen compared to the non-chosen target location (Kim and Basso, 2008, 2010; Pastor-Bernier and Cisek, 2011). The final motor command is then an average of these two differentially weighted motor commands. This model can be conceptualized as a race between two accumulators (Schall, 2001, Enachescu et al., 2021), with the eventual chosen target location accumulating at a faster rate compared to the non-chosen target location. Instead of fitting the weights of the chosen and non-chosen target locations, we decided to indirectly estimate them by using the *Fast* and *Slow RT* tuning curves from the *Single Target* trials, respectively (**Figure 2b)**. Previous studies (Pruszynski et al., 2010; Gu et al., 2016, 2018) have linked trial-by-trial EVR magnitude to the ‘readiness’ of the motor system towards a specific target location. Here, we assumed that during *Double Target* trials the motor system reaches towards the more ‘ready’ target location.

### Weighted motor averaging model best explains EVR-related EMG-activity

**Figure 5a** and **b** summarize the four different model predictions (color lines) for both the ΔPD averaging ratio and normalized amplitude fits across the three different *Double Target* angular separation conditions relative to *Single Target* trials. The *winner-takes-all model* predicted no change in either ΔPD (i.e. averaging ratio = 0 a.u.) or amplitude (i.e. normalized amplitude = 1 a.u.). Both the *spatial* and *motor averaging models* predicted complete averaging (averaging ratio = 1 a.u.) for both the 60° and 120° conditions. The key difference between the two models was in the predicted amplitude, where the *spatial averaging model* predicted no change (amplitude = 1 a.u.), while the *motor averaging model* predicted a systematic decrease (amplitude < 1 a.u.). Finally, the *weighted motor averaging model* predicted both a partial averaging (0 < averaging ratio < 1) and a decrease in amplitude. The extent of these changes depended on each participant’s own *Fast* and *Slow RT* fits.

**Figure 5:**
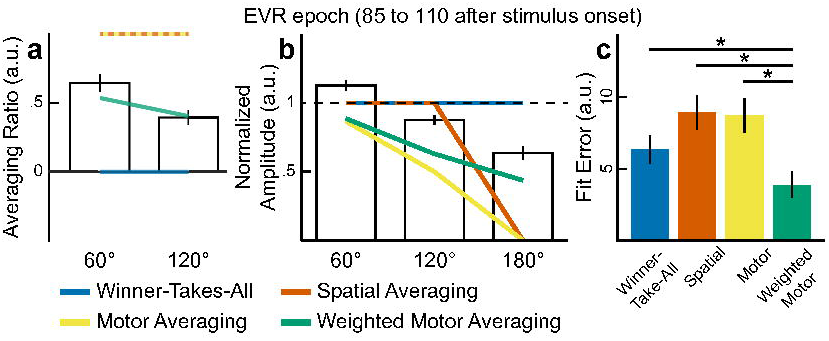
Comparisons of model predictions and observed group data for *Double Target* trial fits. **a, b.** The model predictions (colored lines, see legend for color coding) overlaid over the observed mean ± SEM group data (open black bars) for EMG-activity during the EVR-epoch (85 to 125 ms after stimuli onset) for both the averaging ratio (**a**) and amplitude (**b**). **c.** The mean ± SEM group model fit errors for the four different models.

**Figure 5a** and **b** also show our observed group data (open bars) plotted against the predictions from the four models during the EVR-epoch. Note only the *weighted motor averaging model* (green lines) captured both the systematic decrease in averaging ratio and amplitude that was in the observed data. Since the parameters of all four models were derived from each participant’s own *Single Target* trials and contained no free parameters, we can directly compare the four different models. **Figure 5c** illustrates the mean ± SEM of the fit error between the observed and predicted fits across the participants. We found that the *weighted motor averaging model* best predicted the observed tuning curves compared to the other three models during the EVR-epoch (repeated measures 1-way ANOVA, *F*_(3,30)_ = 7.7, *P* < 10^-3^, post-hoc paired *t*-test, *t*_(10)_ = 3.6, 4.1, and 4.8, *P* = 0.005, 0.002, and 0.0001, compared to the *winner-takes-all, spatial*, and *motor averaging models*, respectively).

### Winner-takes-all model best explains MOV-related EMG-activity

Up to this point, we have only examined the initial wave of EMG-activity time-locked to the onset of the two visual targets (i.e. during the EVR-epoch). Are there also signatures of averaging in the tuning of EMG-activity associated with movement onset (MOV-epoch) in the *Double Target* trials? We therefore examined EMG activity during the MOV-epoch, i.e. mean EMG-activity −20 to 20 ms around reach onset.

**Figure 6a** shows the EMG-activity during the MOV-epoch for individual trials for our exemplar participant, centered at the chosen target direction and split by the direction of the non-chosen target and the three target separations. On top the cosine tuning curves are shown. For this subject the amplitudes do not differ between target separations or non-chosen target direction. However, small shifts in preferred direction, away from the non-chosen target, can be observed for the 60° and 120° target separation.

**Figure 6:**
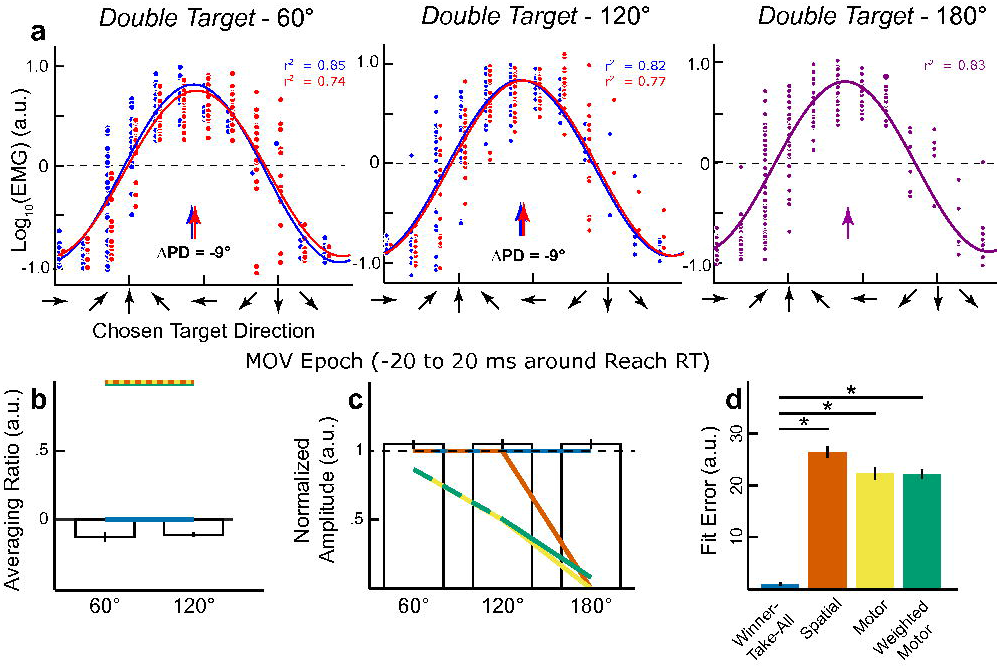
EMG-activity and tuning properties in the MOV-epoch (−20 to 20 ms around reach RT). **a.** Fits for 60°, 120° and 180° conditions of the *Double Target* trials with all data aligned to the chosen target direction from the representative participant. For both the 60° and 120° conditions, the trials were sorted based on whether the non-chosen target was either CW (blue) or CCW (red) of the chosen target direction. **b, c.** The model predictions overlaid over the observed mean ± SEM group data (open black bars) for EMG-activity during the MOV-epoch for both the averaging ratio (**b**) and amplitude (**c**). **d.** The mean ± SEM group model fit errors for the four different models. * *P* < 0.0083.

**Figure 6b** and **c** show both the averaging ratio and amplitude across participants, based on fits to the EMG-activity during the MOV-epoch. Unlike the EVR-epoch, the *winner-takes-all model* best predicted EMG-activity around reach onset (**Fig. 6d**, repeated measures 1-way ANOVA, *F*_(3,30)_ = 348.8, *P* < 10^-22^, post-hoc paired *t*-test, *t*_(10)_ = 28.6, 21.8, 29.0, all *P* < 10^-9^, compared to the *spatial, motor*, and *weighted motor averaging models*, respectively). Although the *winner-takes-all* model provides the best explanation for our MOV-epoch data, we still observed an influence of the non-chosen target location with an averaging ratio shifting in the opposite direction, suggesting a *repulsion* from the non-chosen target location (averaging ratio = −0.13 ± 0.03 and −0.11 ± 0.01 a.u., for 60° and 120° *Double Target* trials, respectively, one sample *t*-test against zero, *t*_(10)_ = −4.2 and −11.7, both *P* < 0.05, **Fig. 6b**). However, in the next section we will argue that this is not a genuine repulsion from the non-chosen target, but rather compensation for the earlier attraction by the non-chosen target in the EVR-epoch.

### Early kinematics show attraction to the non-chosen target location

Having established an opposite influence of the non-chosen target on the tuning of EMG-activity during the EVR and MOV-epoch, we next determined whether the brief burst of muscle recruitment during the EVR-interval carried any behavioral consequences. **Figure 7a** shows the representative participant’s initial reach error (i.e. the difference between the chosen target location and the initial reach direction at the time of reach onset) for both *Single* and *Double Target* trials. For the *Single Target* trials the distribution of initial reach direction is closely centered on the actual target direction. However, for the *Double Target* trials the distributions of initial reach direction are clearly shifted toward the non-chosen target. **Figure 7b** shows the median initial reach error direction, averaged across participants. This initial reach error differed significantly from zero for both target separations (Initial Reach Error = 15.2° ± 4.3° and 13.9° ± 5.5°, paired *t*-test, *t*_(10)_ = −11.7 and −8.4, both *P* < 10^-5^, respectively). These initial reach errors indicate an early attraction toward the non-chosen target, which is consistent with the averaging of EMG-activity during the EVR-interval. Following this averaging during the EVR-interval, we subsequently observed an opposite effect in the tuning of the EMG-activity in the MOV-epoch for the *Double Target* trials. This is highlighted by a significant negative averaging ratio (**Figure 6b**). We surmise that this opposing effect corresponds to compensatory muscular activity that corrects for the initial attraction of the arm toward the non-chosen target, bowing the arm back toward the chosen target location (**Figure 7c).**

**Figure 7:**
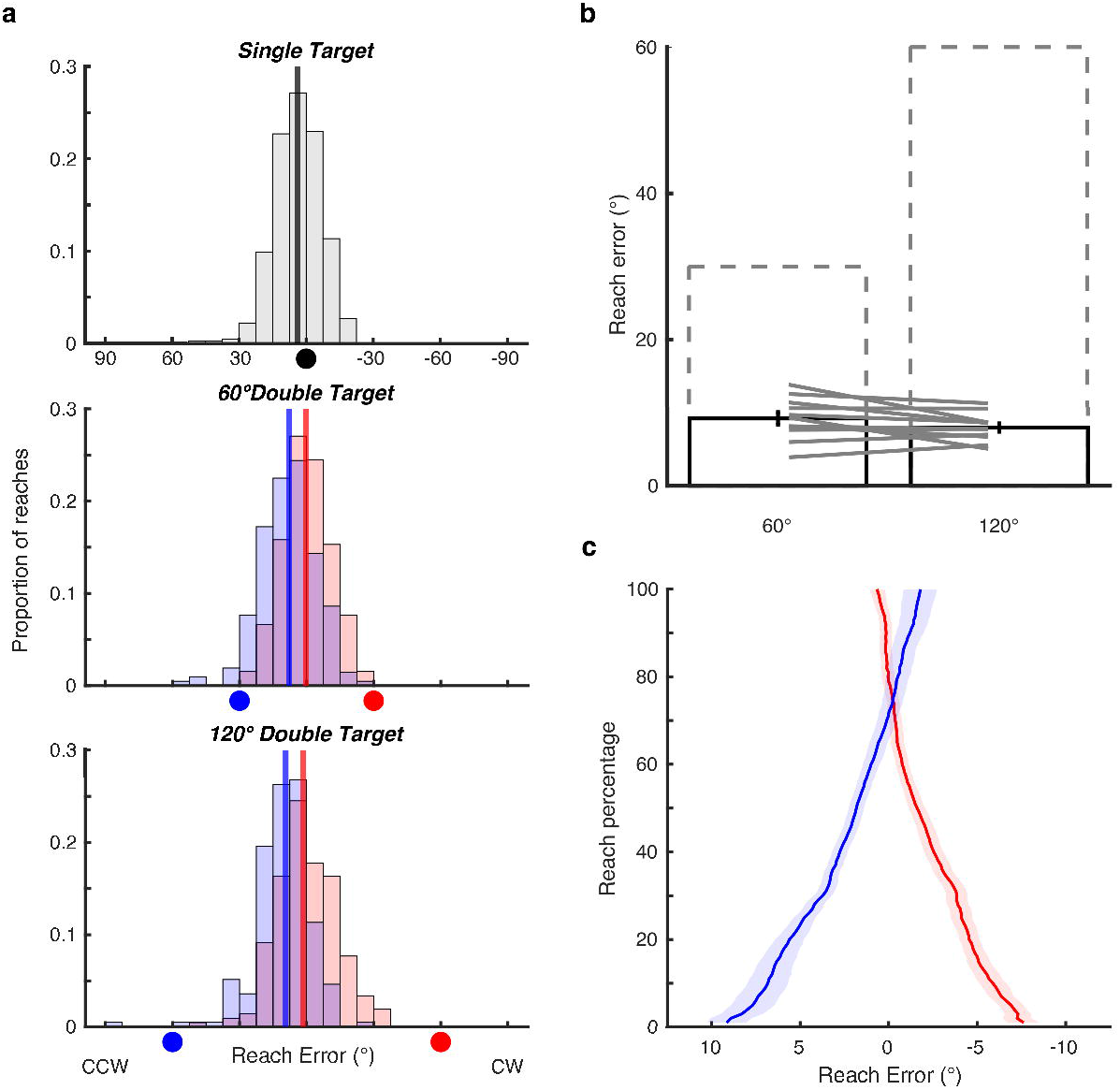
Systematic repulsion away from the non-chosen target direction at the time of reach RT. **a.** Histogram of reach error direction, relative to the chosen target direction, at the time of reach RT for the representative participant during the experiment. For *Double Target* trials, the location of the non-chosen target direction is shown as colored circles along the *x*-axis. Vertical lines represent the median reach errors. **b.** Mean ±SEM of difference in median reach error between CW and CCW during *Double Target* trials. Dashed boxes represent full averaging, i.e. predictions from model 2 and 3. **c.** Initial reach errors converge to the target direction while the reach unfolds and the reach percentage (RP) increases. RP = 0% corresponds to hand speed above 2cm/s and RP = 100% to covering the target distance. Across subjects and conditions this corresponds to a time window of 270 ±23ms.

## DISCUSSION

Contemporary theories of decision-making posit that multiple potential motor plans compete for selection (Cisek, 2007; Schall, 2001). Behavioral and neurophysiological results have shown such competition within the oculomotor system (Chou et al., 1999; Coren and Hoenig, 1971; Findlay, 1982; Ottes et al., 1984; Walker et al., 1997; Bhutani et al., 2012), but it is unclear whether such results generalize to reaching. Here, by measuring upper limb EMG during a reaching task that demands an immediate response, we demonstrate that the non-chosen target influences the earliest wave of muscle recruitment following target onset. Such evidence is apparent on single trials, implicating biased competition between the chosen and non-chosen target within upstream premotor areas soon after target appearance. This initial biased motor averaging affected the initial direction, but subsequently gave way to a goal-directed motor command that bowed the arm back onto a trajectory directed toward the chosen target.

### The EVR is a trial-by-trial weighted average of motor commands

We tested different models of how the brain could have integrated the two visual targets and found that a *weighted-motor-averaging* model best explained partial averaging during the EVR-epoch. Such weighted-motor-averaging is apparent on a single trial and inconsistent with a recent interpretation that the apparent encoding of multiple alternatives in premotor cortex is caused by averaging of different alternatives across multiple trials (Dekleva et al., 2018). Instead, our result is consistent with contemporary theories for a deliberation process between multiple motor plans (Cisek, 2007; Schall, 2001; Enachescu et al., 2021). For example, previous neurophysiological studies have shown that experimentally manipulating the decision variable, via either target uncertainty (Basso and Wurtz, 1997; Dorris and Munoz, 1998), target expectation (Basso and Wurtz, 1998; Bichot et al., 1996), or reward expectation (Pastor-Bernier and Cisek, 2011; Rezvani and Corneil, 2008), modulates the neural representation of the competing motor plans. Similarly, signatures of an evolving decision variable during deliberation have been shown in the long latency reflex, when participants must indicate the direction of a random-dot motion stimulus (Selen et al., 2012). Here, we exploited the fact that participants randomly choose one of two visual targets and demonstrated post-hoc that the averaged EVR was biased towards the chosen target. This suggests that either fluctuations along the sensorimotor pathway (Faisal et al., 2008; Siegel et al., 2015) or idiosyncratic preferences based on previous choices (Urai et al., 2019) biased both the initial EVR and the ultimate choice. Interestingly, the influence of idiosyncratic preferences on the representation of alternatives was also reported in premotor cortex (Dekleva et al., 2018).

While the weighted-motor-averaging model best explained the EVR, the fits were imperfect (**Figure 5**). This is likely due to the arbitrary weighting of the EVR strength for the chosen vs. non-chosen target, exploiting the inverse relationship between EVR magnitude and reach RT (Gu et al., 2016; Pruszynski et al., 2010; Wood et al., 2015). We assume an independent race between the motor programs to each of the two targets, with the one proceeding faster (drawn from the shorter-than-average subset, having a larger EVR) ‘winning’ over the non-chosen alternative (Rowe et al., 2010). This process is admittedly coarse, as it remains unknown what the RT and EVR of the non-chosen alternative would have been. Regardless, only the weighted-motor-averaging model captured the influence of the non-chosen target on both the EVR tuning and magnitude (**Figure 5**); hence this model best captures the essence, if not the magnitude, of the interaction between competing motor plans.

A *weighted-spatial-averaging* model, an extended version of **Model 2,** was not explicitly evaluated. If we would allow the averaged target location to be somewhere between the presented targets, instead of in the middle, the pre-motor circuitry would receive a ‘go here’ signal that could change the shift of the tuning curve, but would not influence the amplitude of the tuning. In contrast, our data show a systematic decrease in amplitude for the *Dual Target* conditions, which can only be captured by the *weighted-motor-averaging* model.

### Influence of task design on the EVR

Our results illustrate that different stages of decision-making influence distinct EMG epochs in the motor periphery and thus suggest an influence of task design or stimulus properties on these epochs. Indeed, the EVR is muted (Wood et al., 2015) or abolished (Pruszynski et al., 2010) when a delay is imposed between stimulus presentation and movement onset. Furthermore, the EVR is augmented when targets are temporally predictable (Kozak et al., 2020; Contemori et al., 2021a). Given this, the EVR may be negligible or absent in delayed response tasks (Cisek and Kalaska, 2005; Dekleva et al., 2018; Thura and Cisek, 2014), or in *‘go-before-you-know*’ tasks introducing a delay between presentation of alternatives and initiation of the reach. When an immediate response is required in the *‘go-before-you-know’* paradigm, the intermediate reaching movements skew toward the more salient stimulus (Wood et al., 2011), paralleling the observation of earlier and larger-magnitude EVRs evoked by high-contrast (Wood et al., 2015; Kozak and Corneil, 2021) or low-spatial frequency stimuli (Kozak et al., 2019).

The instruction to move rapidly reduces the production of intermediate reaches, possibly due to adopting a control policy that maximizes task success (Wong and Haith, 2017). While the impact of velocity instructions on the EVR is unknown, previous results suggest that the magnitude, not timing, of the EVR would be modulated by changing control policy (Gu et al., 2018, 2016). Furthermore, the EVR’s short-latency makes the establishment of a control policy after target presentation unlikely, but suggests a task dependent, preset control policy implementing task instructions affecting relevant motor circuitry (Scott, 2016; Contemori et al., 2022). Recently, Enachescu et al. (2021) provided a dynamic neural field model connected with stochastic optimal feedback controllers. This model executes a weighted average of a continuum of control policies for all possible reach directions, where competition and weighing of control policies continues as the reach unfolds, consistent with the present findings.

### Kinetic consequences of the EVR

EMG recordings permit the resolution of a decision-making dynamic at a level that would be difficult, if not impossible, to resolve based on kinematics alone. For example, while EMG-activity during the EVR was biased *toward* the non-chosen target, EMG-activity during the MOV-interval was biased *away* from the non-chosen target (**Fig. 6**). At first glance, opposite directions of EMG recruitment in these intervals seems paradoxical. The forces consequent to the brief and smaller-magnitude EVR are undoubtedly less than those developed closer to the time of reach initiation. However, the EVR has behavioral consequences, generating small forces toward a stimulus (Gu et al., 2016). We show that forces from the averaged EVR bias the initial reach toward the non-chosen target, but subsequent EMG compensates for their trial-specific kinematic consequences. This suggests that voluntary control mechanisms are rapidly informed about trial-specific kinematic consequences of the averaged EVR, using this information for feedforward adjustments of the voluntary EMG-activity. These adjustments occur within 100ms after the onset of EVR and are unlikely to be driven by visual feedback of the cursor. A similar fast mechanism has been reported for stretch reflexes (Pruszynski et al., 2009). If the background load to a muscle increases, the mono-synaptic short latency reflex increases, but adjustments in later phases, as quick as 45 ms after perturbation onset, already compensate for the stronger adjustment in the first phase.

### A shared neural substrate with the saccadic system

Our task incorporated many task features used to elicit saccadic averaging (Chou et al., 1999; He and Kowler, 1989), including the requirement for an immediate response. Most experiments on saccadic averaging have not been designed to dissociate between averaging at the spatial or motor level. However, by contrasting two task instructions (“look at the last presented target” vs “look at the targets in order of presentation”) in a double-step paradigm, Bhutani et al., (2012) provided evidence that saccadic averaging also takes place at the level of the motor plan. Saccade kinematics offer a straightforward readout of the temporal evolution of decision-making, paralleling our observations for EVRs. For example, the transition from an averaged to a goal-directed command between the EVR and MOV-epoch resembles the observation that averaging is strongest for short-latency saccades (Chou et al., 1999; Walker et al., 1997). Further, EVR averaging diminishes with increasing angular target separation, resembling observations for saccadic averaging (Chou et al., 1999; Vokoun et al., 2014). Saccadic averaging has been related to the initial representation and subsequent resolution of competing saccade plans within superior colliculus (Edelman and Keller, 1998; Port and Wurtz, 2003; Vokoun et al., 2014). Superior colliculus is also a potential substrate for the EVR via the tecto-reticulo-spinal pathway (Corneil and Munoz, 2014; Glover and Baker, 2019; Gu et al., 2016; Pruszynski et al., 2010; Contemori et al., 2021a, Kozak and Corneil, 2021; Kozak et al., 2019). Thus, circumstantial evidence suggests that saccadic averaging and EVR averaging on upper limb muscles may have a common collicular substrate. This subcortical substrate for the deliberation process would agree with findings that M1 and PMd are mainly involved in commitment to a choice (Derosiere et al., 2019; Thura and Cisek, 2020), but not the competition between alternatives. Future neurophysiological experiments should investigate the causal structure between weighted averaging of the EVR and the commitment to a single goal-directed reach.

In summary, we examined neuromuscular activity during a free-choice reaching task to two targets. We found that, similar to saccadic averaging, the earliest motor command in the reaching system attests to a still-unresolved competition between multiple distinct motor plans. However, this competition is rapidly resolved and by the time of movement onset the motor system generates a goal directed reach movement that compensates for the averaging observed in the early trajectory.

## Acknowledgement

This work was supported by operating grants from the Natural Sciences and Engineering Research Council of Canada (NSERC) to BDC [RGPIN-311680], the Canadian Institutes of Health Research (CIHR) to BDC [MOP-93796], a Vici grant from the Netherlands Organization for Scientific Research to WPM [453-11-001].

## AUTHOR CONTRIBUTIONS

Conceptualization – LPS, BDC, and WPM; Methodology – LPS; Investigation – LPS; Writing, Original Draft – LPS; Writing, Review and Editing – LPS, BDC and WPM; Funding Acquisition – WPM; Resources – LPS and WPM.

## DECLARATION OF INTERESTS

The authors declare no competing interesting.

## Notes

### Competing Interest Statement

The authors have declared no competing interest.

### Summary of Updates

Updated the paper based on reviews. No fundamental changes, just further clarification, improved writing.

